# Biosynthesis of *Securinega* alkaloids via a cyanogenic glucoside-like pathway

**DOI:** 10.64898/2026.10.14.759206

**Authors:** Zhouqian Jiang, Lachlan J. N. Waddell, Inesh Amarnath, Caitlin A. Hancock, Owen R. Hughes, Gideon Grogan, William P. Unsworth, Benjamin R. Lichman

## Abstract

The elucidation of biosynthetic pathways towards bioactive compounds in plants is essential for building biotechnological platforms for their scalable synthesis. Securinine and allosecurinine form part of the complement of *Securinega* alkaloids in *Flueggea suffruticosa*, a fundamental herb of Chinese medicine. Despite their potent biological activities and a growing interest in derivatives as pharmaceutical leads, key steps in their biosynthesis remain obscure. Here we report the discovery and reconstitution of the pathway to the core *Securinega* alkaloid precursor menisdaurilide. Using transcriptomics, *Nicotiana benthamiana* transient expression, *in vitro* assays, and pathway reconstitution in *Saccharomyces cerevisiae*, we reveal that three cytochrome P450s and a reductase are required for the biosynthesis of menisdaurilide from L-tyrosine via the cyano-metabolite menisdaurigenin, revealing a surprising convergence with the biosynthesis of cyanogenic glucosides such as dhurrin. The *de novo* biosynthesis of allosecurinine in *N. benthamiana* highlights the potential for preparative-scale heterologous production of *Securinega* alkaloids.

## Introduction

Alkaloids are a large and diverse class of plant natural products characterised by the presence of a basic nitrogen^1^ and often produced for defensive properties including protection against herbivory^2^. They can exhibit potent physiological activity in animals and humans^3^; consequently, many have pharmacological or stimulatory uses, such as morphine, quinine and nicotine^4^.

Securinine (**1**) and its epimer allosecurinine (**2**) are tetracyclic alkaloids derived from the East Asian medicinal plant *Flueggea suffruticosa* (syn *Securinega suffruticosa*)^5,6^. Securinine (**1**) is a γ-aminobutyric acid (GABA_A_) receptor antagonist^7^ and exhibits anticancer^8^, antimicrobial, and anti-inflammatory activities^9^ while allosecurinine (**2**) shows anti-fungal activities^10^. These compounds are part of the *Securinega* alkaloid family, found primarily in the Phyllanthaceae^11–13^. The alkaloids and their derivatives are under investigation for diverse pharmaceutical applications^9,14,15^. Elucidating their origins could enable bio-based technologies for their production.

Unlike many specialised metabolite classes that share unified biosynthetic origins, alkaloids show lineage-specific distribution, emerging from scaffold-forming steps that lack conservation across taxa. Nevertheless, their assembly follows a biochemical logic: a Mannich-like condensation of an amino acid-derived imine electrophile and a corresponding nucleophile^1^. Piperidine alkaloids derived from lysine (**3**) employ Δ^1^-piperideine (**4**) as a cyclic imine electrophile. Securinine (**1**) and allosecurinine (**2**) are derived from lysine (**3**)^16,17^; we recently identified a *F. suffruticosa* enzyme, *Fs*OLADO, which catalyses the conversion of lysine (**3**) to Δ¹-piperideine (**4**)^18^.

The metabolic origin of the nucleophile that contributes to lysine derived alkaloids is highly varied: for example, in lupin alkaloids like sparteine or lupanine the nucleophile is derived from a second Δ^1^-piperideine via tautomerisation^19^. The *Lycopodium* alkaloids (*e.g*. huperzine A) derive their nucleophiles from polyketides^20^, whereas anabasine and related *Nicotiana* alkaloids use a nicotinic acid derived nucleophile activated by glucosylation and reduction^21^. The *Securinega* alkaloids appear to be unique among lysine (or ornithine) derived amino acids in deriving their nucleophilic moiety from a second amino acid, tyrosine (**5**)^16,22–24^.

A breakthrough occurred when Busqué, de March, and colleagues, inspired by the identification of the butenolide menisdaurilide (**6a**) and its epimer aquilegiolide (**6b**) in *Securinega* alkaloid producing plants^25,26^, synthesised allosecurinine (**2**) from menisdaurilide (**6a**)^27^. The key steps of this and related syntheses are Mannich-like reactions in which a menisdaurilide (**6a**) derivative acts as a nucleophile combining with a piperidinium-like electrophile^27–29^.

Recently, we demonstrated that Δ¹-piperideine (**4**) and menisdaurilide (**6a**) can undergo a non-enzymatic Mannich-type reaction in aqueous conditions to form the neosecurinane alkaloids virosine B (**7**), virosine A (**8**) and securinol A (**9**)^30^ (**Fig. 1A**). In parallel, the Han and Kim groups identified a sulfotransferase (*Fs*NSST1) that catalyses the scaffold remodelling of virosine B (**7**) and virosine A (**8**) into allosecurinine (**2**) and securinine (**1**) respectively^31^.

**Figure 1.**
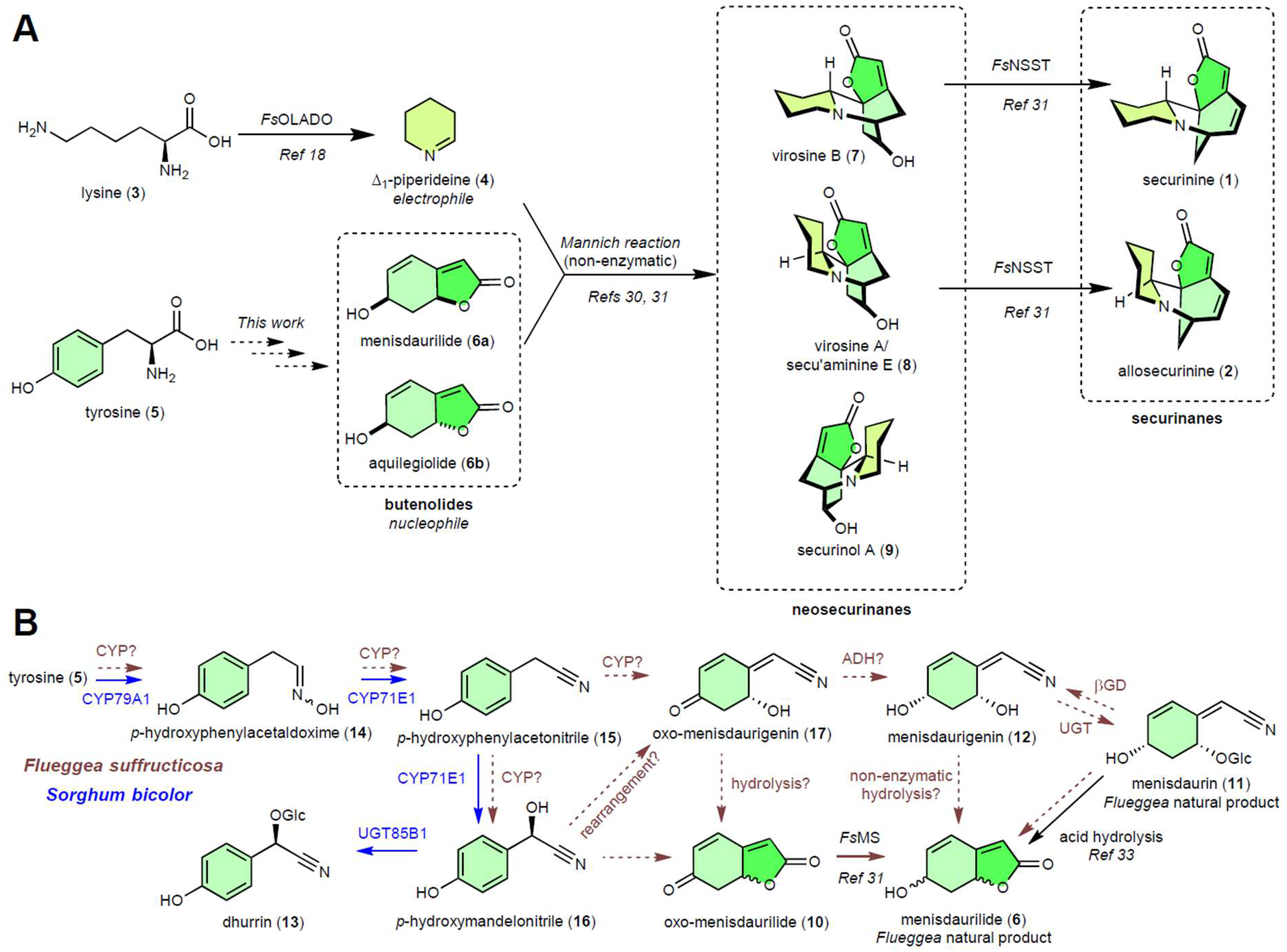
The biosynthesis of *Securinega* alkaloids. (**A**) Proposed biosynthetic pathway of major *Securinega* alkaloids. Dashed lines indicate unresolved steps. (**B**) Proposed *F. suffruticosa* menisdaurilide (**6**) biosynthesis pathway (green) in comparison to *Sorghum bicolor* dhurrin biosynthesis (blue).

With Δ¹-piperideine (**4**) formation^18^, the Mannich reaction^30^ and subsequent scaffold remodelling established^31^, the remaining missing link for the bio-based formation of *Securinega* alkaloids is the biosynthetic route converting tyrosine (**5**) into menisdaurilide (**6a**). Recently a retrosynthesis algorithm predicted the origin of menisdaurilide from tyrosine via 4-hydroxyphenylacetic acid and an unusual *para-*quinone methide intermediate^32^. Furthermore, an NADPH-dependent reductase, menisdaurilide synthase (*Fs*MS), was reported to catalyse the reduction of oxo-menisdaurilide (**10**) to menisdaurilide (**6a**)^31^.

Menisdaurilide (**6a**) forms during the acid hydrolysis of menisdaurin (**11**)^33^, a nitrile glucoside first isolated from *Menispermum dauricum*^34^ and also found in *Flueggea* species^35^. The aglycone of menisdaurin (**11**), menisdaurigenin (**12**), is unstable in water and methanol but can be prepared with enzymatic deglucosylation and extraction into organic solvent^33^. Menisdaurin (**11**) is structurally similar to the tyrosine derived cyanogenic glucoside dhurrin (**13**) and they have been reported to co-occur in some species: their biosynthetic origins may therefore share common features (**Fig. 1B**)^36^.

In this study we set out to discover the biosynthetic and enzymatic origins of menisdaurilide (**6**), as a first step to establishing a platform for alkaloid formation in heterologous systems. First, we hypothesise that menisdaurilide (**6**) formation shares features with cyanogenic glucoside pathways. Using co-expression data, *N. benthamiana* assays, *in vitro* enzyme validation and heterologous pathway reconstruction, we discover and characterise four enzymes (three cytochrome P450s and one reductase) that convert tyrosine (**5**) to menisdaurilide (**6**). We show the pathway proceeds via the reactive intermediate menisdaurigenin (**12**) and shares a close biochemical connection with cyanogenic glucoside formation. Finally, we use this newly identified pathway for heterologous construction of a biosynthetic pathway for *Securinega* alkaloids including allosecurinine (**2**).

## Results

### The menisdaurilide biosynthetic pathway

To identify menisdaurilide (**6**) biosynthetic genes, we first developed a pathway hypothesis inspired by the previously observed acid-catalysed conversion of menisdaurin (**11**) to menisdaurilide (**6**)^26^, the detection of menisdaurin (**11**) in plants from *Securinega* alkaloid producing genera^25,35^, and the potential biosynthetic connection between cyanogenic glucosides like dhurrin (**13**) and nitrile glucosides^36^. We proposed that two or three cytochrome P450s and a reductase were involved in the pathway from tyrosine (**5**) to menisdaurilide (**6**) via the intermediate menisdaurigenin (**12**) (**Fig. 1B**). The putative pathway follows dhurrin (**13**) biosynthesis through *p*-hydroxyphenylacetaldoxime (**14**) and *p*-hydroxyphenylacetonitrile (**15**), where it bifurcates, not forming *p*-hydroxymandelonitrile (**16**) but instead oxo-menisdaurigenin (**17**). This proposed intermediate undergoes reduction to menisdaurigenin (**12**) and hydrolysis to yield menisdaurilide (**6**).

As is typical in plant biosynthetic pathways, we expected that these genes would co-express with *Fs*OLADO, the enzyme that provides Δ¹-piperideine (**4**) to *Securinega* alkaloids^18^. Using gene expression data, derived from fifteen different tissues and developmental stages^18^, we identified multiple genes with Pearson correlation coefficient greater than 0.75 relative to *Fs*OLADO, including cytochrome P450s and reductases, as well as UDP-glucosyltransferases (UGTs) and β-glucosidases, which we hypothesised may be involved in the interconversion of menisdaurigenin (**12**) and menisdaurin (**11**) (**Table S1**).

Ten candidate genes were cloned from *F. suffruticosa* and inserted into expression vectors for *Agrobacterium*-mediated transient expression in *Nicotiana benthamiana* (**Table S2**). For gene24453, encoding a cytochrome P450, we identified two isoforms (v1/v2) sharing 90% sequence identity (**Fig. S1A**). Co-infiltration of all ten candidate genes led to the *de novo* production of menisdaurilide (**6a**) and aquilegiolide (**6b**) (**Fig. 2A** and **B**). Gene dropout deconvolution revealed that cytochrome P450s gene2189 and gene7595, as well as the reductase gene1004 were necessary for menisdaurilide (**6**) production (**Fig. 2C**). However, co-expression of these three genes was not sufficient to induce menisdaurilide (**6**) production (**Fig. 2D**).

**Figure 2.**
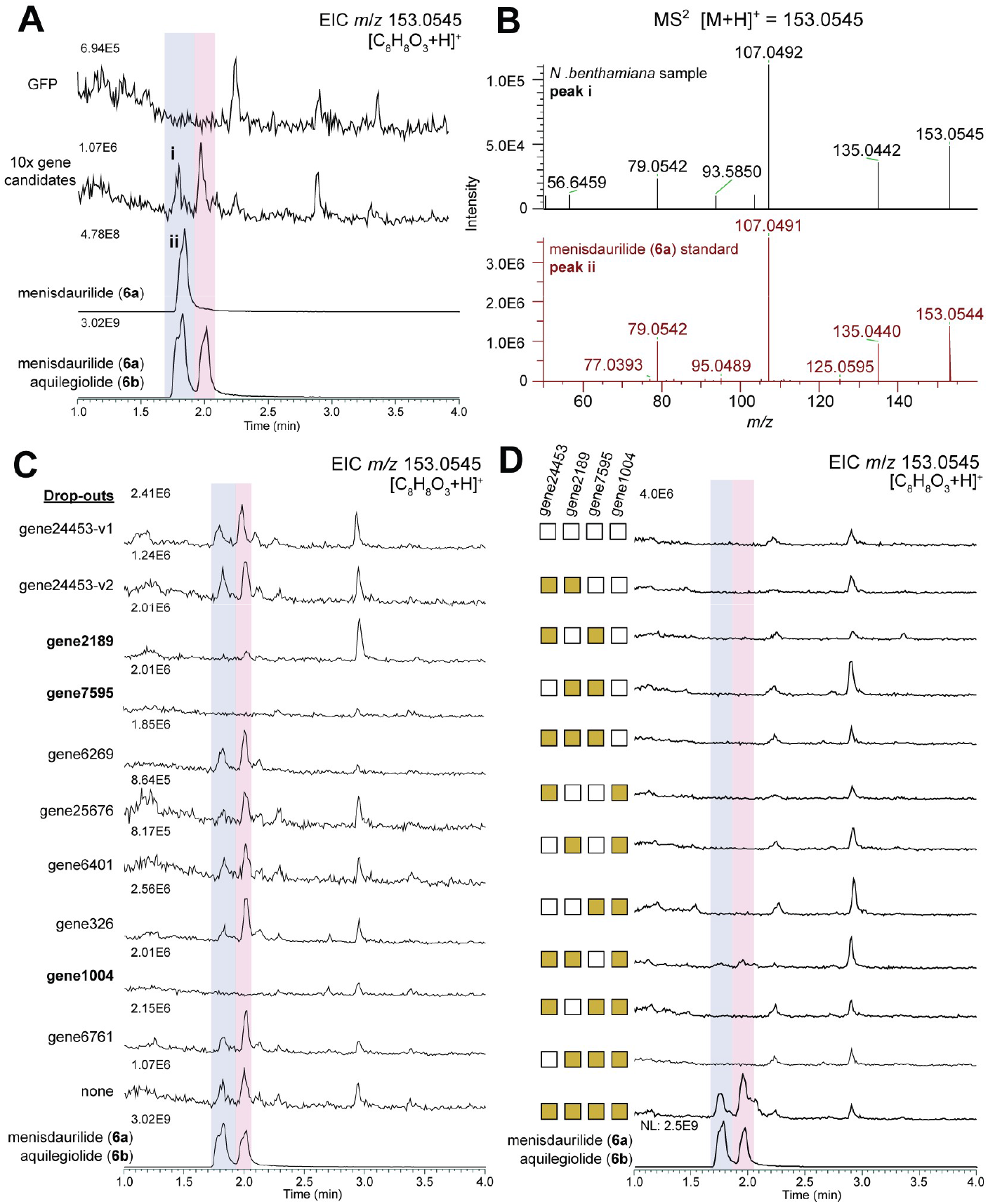
The identification of menisdaurilide biosynthesis genes. (**A**) LC-MS extracted ion chromatograms (EICs) for menisdaurilide (**6**) [M+H]^+^ showing menisdaurilide (**6**) production in *N. benthamiana* leaves co-expressing ten candidate genes compared to GFP control and standards. Peaks used in MS^2^ confirmation (see panel **B**) are labelled. (**B**) MS² fragmentation spectra of *m*/*z* 153.0545 peaks from standard and sample labelled in panel **A**. (**C**) EICs showing menisdaurilide (**6**) content in single gene drop-out experiments in *N. benthamiana* leaves. Samples were infiltrated with all ten candidate genes excluding those labelled as drop-outs. (**D**) EICs showing menisdaurilide (**6**) accumulation in *N. benthamiana* leaves with different gene combinations infiltrated. Yellow boxes show the genes infiltrated in each sample. Gene24453-v1 was used here, see **Fig. S1B** for gene24453-v2 activity.

We predicted that, due to their high sequence similarity, the cytochrome P450s gene24453-v1 and gene24453-v2 were functionally redundant, leading to failure of the drop-out experiment to reveal their involvement. Co-expression of gene24453-v1 (or gene24453-v2) alongside gene2189, gene7595, and gene1004, revealed these four genes to be sufficient for the *de novo* production of menisdaurilide (**6a**) and aquilegiolide (**6b**) in *N. benthamiana* (**Fig. 2D**, **Fig. S1B**). The newly identified cytochrome P450s were named *Fs*CYP79DC12/13 (gene24453-v1/v2), *Fs*CYP81BD45 (gene2189), *Fs*CYP92B120 (gene7595).

*Fs*CYP79DC12 and *Fs*CYP81BD45 are adjacent on the *F. suffruticosa* genome assembly, and *Fs*CYP92B120 is ∼7 Mbp away on the same chromosome (**Figs. S2**)^31^. The same genomic organisation can be found for the orthologs on the genome of the alkaloid producing *Flueggea virosa*^37^. During our investigation the reductase gene1004 was reported and characterised as menisdaurilide synthase (*Fs*MS)^31^ and is located on a different chromosome to the cytochrome P450s.

### *Fs*CYP79DC12/13 are tyrosine *N*-monooxygenases

We next elucidated each step from tyrosine (**5**) to menisdaurilide (**6**). Expression of individual cytochromes P450 in *N. benthamiana* did not lead to the accumulation of tyrosine derived *p*-hydroxyphenylacetaldoximes (**14**) (**Fig. S3**). However, the *O*-glucoside of *p*-hydroxyphenylacetaldoxime (**18**) was observed upon expression of *Fs*CYP79DC12 or *Fs*CYP79DC13 (**Figs. 3A, 3B,** and **S4A**). This is a likely product of endogenous *N. benthamiana* glucosyltransferase activity on *p*-hydroxyphenylacetaldoxime (**14**). To confirm the function of *Fs*CYP79DC12 and *Fs*CYP79DC13 as tyrosine *N*-monooxygenases, we performed *in vitro* assays using yeast-derived microsomes. Microsomes expressing either *Fs*CYP79DC12 or *Fs*CYP79DC13 catalysed the conversion of tyrosine (**5**) to *p*-hydroxyphenylacetaldoxime (**14**) (**Fig. 3C**). We also identified a *F. suffruticosa* glycosyltransferase (*Fs*UGT85A242, gene25676) that was able to catalyse the *O*-glucosylation of the aldoxime moiety of *p*-hydroxyphenylacetaldoximes (**14**) to form the corresponding aldoxime glucoside (**19**) (**Figs. 3A, 3D, S4B,** and **S5**).

**Figure 3.**
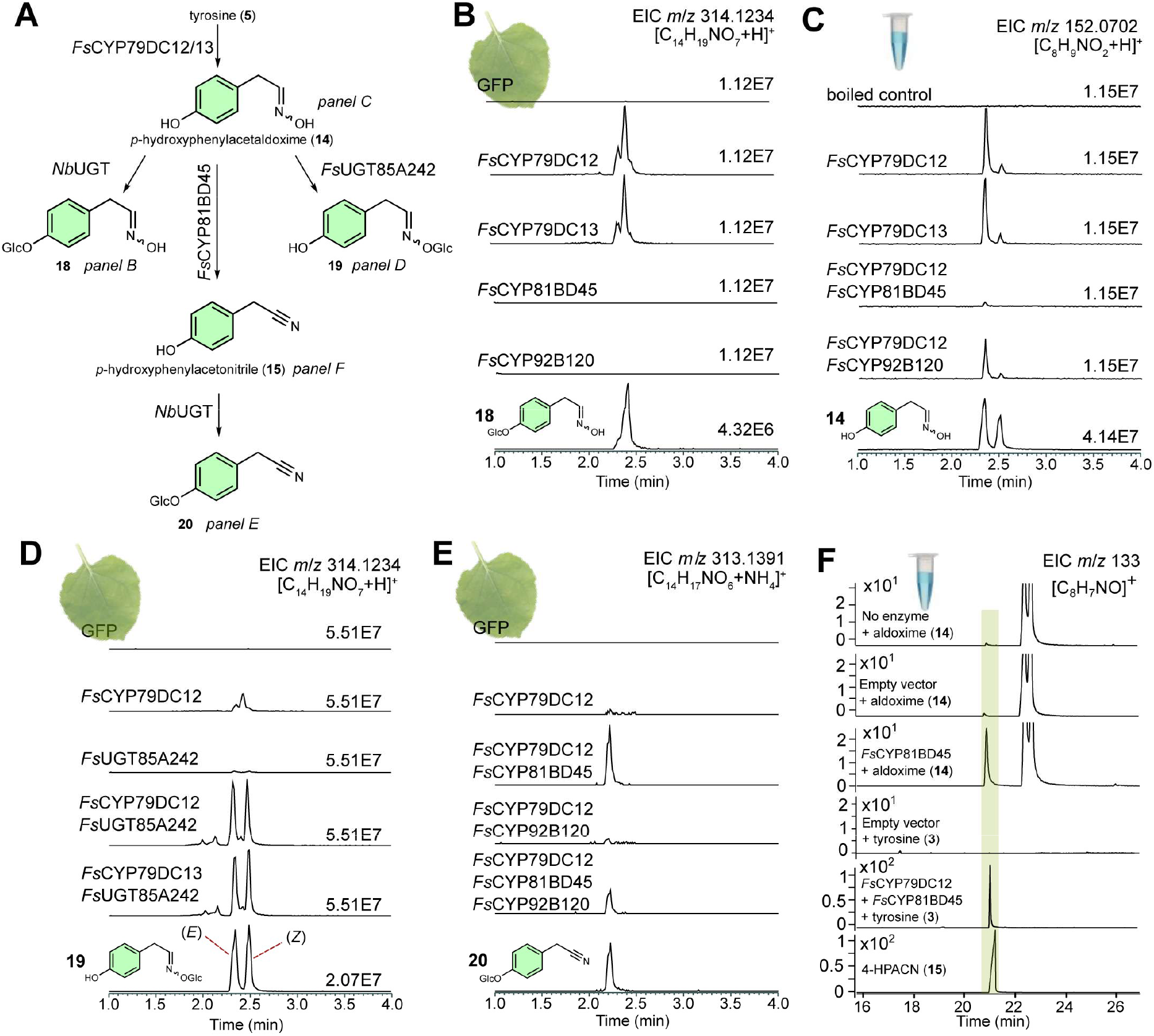
Biosynthesis of 4-hydroxyphenylacetonitrile from tyrosine. (**A**) Overview of the conversions shown in the figure. *Nb*UGT = refers to native *N. benthamiana* glucosyltransferase activity. (**B**) *p*-Hydroxyphenylacetaldoxime phenolic *O*-glucoside (**18**) accumulation in *N. benthamiana* expressing cytochrome P450s. See **Fig. S4A** for MS^2^ characterisation. (**C**) *In vitro* activity of cytochrome P450s on tyrosine (**5**) leading to *p*-hydroxyphenylacetaldoxime (**14**) formation. (**D**) Aldoxime glucosides (**19**) accumulation in *N. bethamiana* with *Fs*CYP79DC12/13 and *Fs*UGT85A242. See **Fig. S4B** for MS^2^ characterisation and **Fig. S5** for *in vitro* confirmation. (**E**) *p*-Hydroxylphenylacetonitrile *O*-glucoside (**20**) accumulation in *N. benthamiana* by *Fs*CYP79DC12 and *Fs*CYP81BD45. See **Fig. S7** for MS^2^ characterisation. For panels **B**-**E** chromatograms are LC-MS EICs with *m/z* as shown on top right of each panel. Values on traces are y-axis normalisation factors. (**F**) GC-MS EIC chromatogram showing *in vitro* conversion of tyrosine (**5**) or *p*-hydroxyphenylacetaldoxime (**14**) to 4-hydroxylphenylacetonitrile (**15**) by *Fs*CYP79DC12 and *Fs*CYP81BD45. See **Fig. S8** for EI fragmentation spectra of **15**.

### *Fs*CYP81BD45 is a 4-hydroxyphenylacetaldoxime dehydratase

Following the characterisation of the first step catalysed by *Fs*CYP79DC12 or *Fs*CYP79DC13, we investigated the functions of the remaining cytochrome P450s. Co-expression of *Fs*CYP81BD45 with *Fs*CYP79DC12 in *N. benthamiana* resulted in a decrease of the *p*-hydroxyphenylacetaldoxime phenolic *O*-glucoside (**18**) relative to the *Fs*CYP79DC12 only sample (**Figs. S6**), and led to the formation of 4-hydroxyphenylacetonitrile *O*-glucoside (**20**) (**Fig. 3E, Fig. S7**). Similarly, combining *Fs*CYP81BD45 and *Fs*CYP79DC12 *in vitro* with tyrosine led to a decrease in aldoxime (**14**) relative to *Fs*CYP79DC12 alone, suggesting that they are substrates for *Fs*CYP81BD45 (**Fig. 3C**). This was confirmed by microsome assays in which incubation of *Fs*CYP81BD45 with aldoximes (**14**) led to the formation of 4-hydroxyphenylacetonitrile (**15**), corresponding to *Fs*CYP81BD45 catalysing the dehydration of 4-hydroxyphenylacetaldoxime (**14**) (**Fig. 3F, Fig. S8**). Notably, when including *Fs*CYP92B120 with *Fs*CYP79DC12 and *Fs*CYP81BD45 in *N. benthamiana*, the yield of **20** decreased, supporting the downstream role of *Fs*CYP92B120 (**Fig. 3E**). This also suggested a sequential reaction order of *Fs*CYP79DC12, *Fs*CYP81BD45, *Fs*CYP92B120, and *Fs*MS.

### Menisdaurilide formation from 4-hydroxyphenylacetonitrile catalysed by *Fs*CYP92B120 and *Fs*MS

To characterise the function of *Fs*CYP92B120, 4-hydroxyphenylacetonitrile (**15**) was fed to *N. benthamiana* leaves expressing *Fs*CYP92B120. This resulted in the production of multiple new peaks with *m/z* 155.0702 [C_8_H_10_O_3_+H]^+^ or *m/z* 154.0860 [C_8_H_11_NO_2_+H]^+^ (**Fig. 4A**, **4B** and **4C**). The *m/z* 155 peaks were determined to be dihydromenisdaurilide (**21a**) and dihydroaquilegiolide (**21b**) (**Fig. 4B, Fig. S9A**). Dihydromenisdaurilide (**21a**) is a natural product previously identified in *Phyllanthus urinaria*^38^. The *m/z* 154 peaks corresponded to *E*-stereoisomers of menisdaurin E ((*E*)-**22a** and (*E*)-**22b**) (**Fig. 4C**, **Fig. S9B**). A Z-stereoisomer of menisdaurin E ((*Z*)-**22**) is a natural product identified in *Bruguiera gymnorrhiza*^39^. The results from *Fs*CYP92B120 and 4-hydroxyphenylacetonitrile (**15**) infiltration were unanticipated as we were expecting an oxidation (-2) or hydroxylation (+16) product of **18** but instead menisdaurin E (**22**) was observed, as a mixture of *E-* and Z-isomers, alongside **15** having undergone a net hydration and reduction (+20).

**Figure 4.**
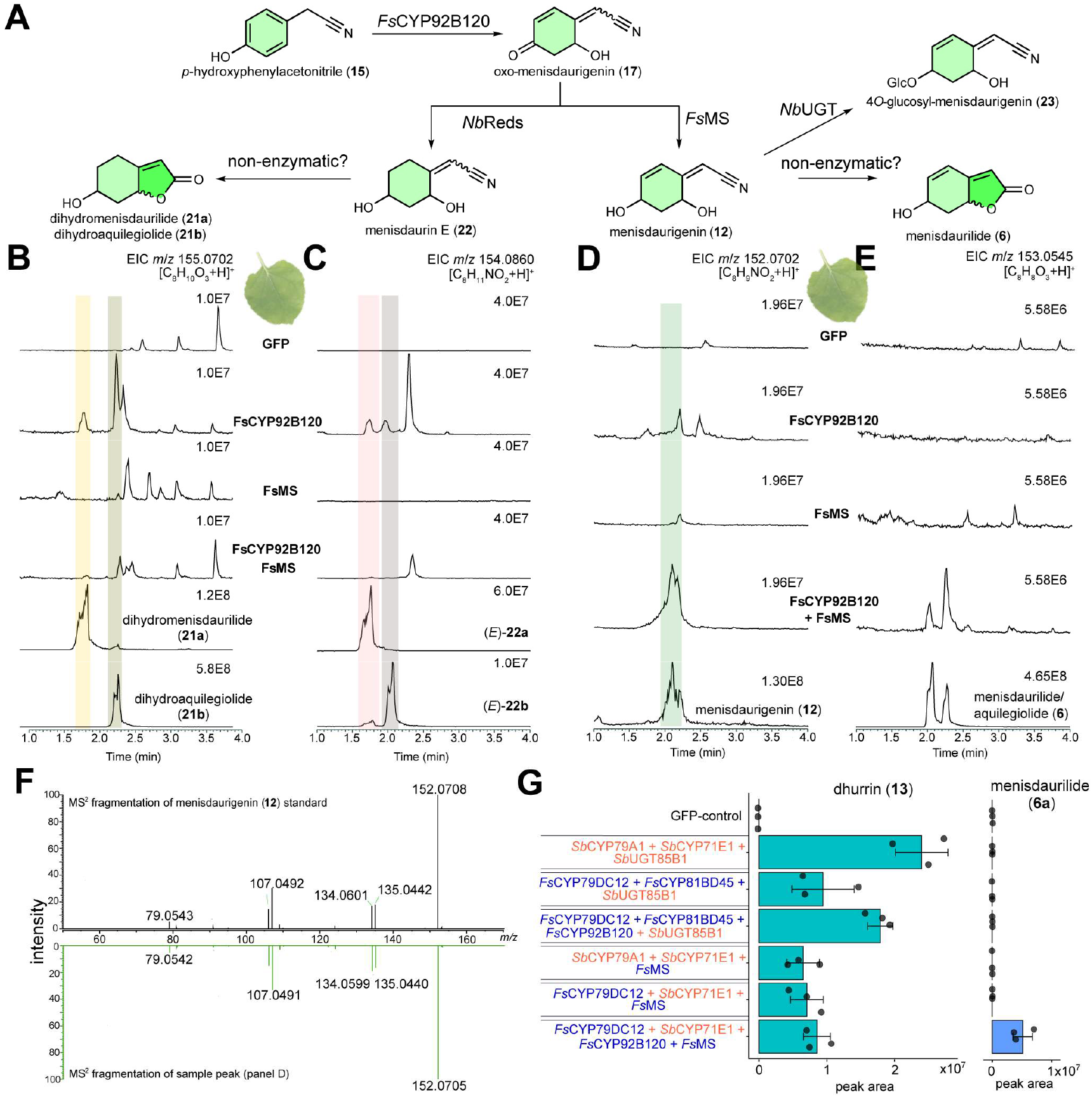
Formation of menisdaurilide from 4-hydroxyphenylacetonitrile. (**A**) The proposed biosynthetic pathway from 4-hydroxyphenylacetonitrile (**15**) to menisdaurilide (**6**) including intermediates and shunt products. *Nb*Reds and *Nb*UGTs refer to proposed *N. benthamiana* reductases and glucosyltransferases respectively. See **Fig. S10** for details of accumulation of **23**. (**B**-**E**) EICs of extracts of *N. benthamiana* leaves fed with 4-hydroxyphenylacetonitrile (**15**) and expressing *Fs*CYP92B120 and/or *Fs*MS: showing dihydromenisdaurilide (**21a**) and dihydroaquilegiolide (**21b**) (panel **B**), menisdaurin E (**22**) (panel **C**), menisdaurigenin (**12**) (panel **D**) and menisdaurilide (**6**) (panel **E**). See **Fig. S9** for further validation of **21** and **22** peak identity. (**F**) MS² validation of menisdaurigenin (**12**) in sample and standard. (**G**) Dhurrin (**13**) and menisdaurilide (**6a**) accumulation in pathway gene substitution experiments in *N. benthamiana*.

When *Fs*CYP92B120 was co-expressed in *N. benthamiana* with the reductase *Fs*MS and supplemented with 4-hydroxyphenylacetonitrile (**15**), the abundance of dihydromenisdaurilide (**21**) and menisdaurin E stereoisomers (**22**) and decreased (**Fig. 4B** and **4C**). Alongside this, there was formation of menisdaurigenin (**12**) (**Fig. 4D** and **4F**), menisdaurilide (**6**) (**Fig. 4E**), and an unknown compound (**23**) (**Fig. S10**). Based on the peak’s *m/z* 314.1234 (matching [C_14_H_19_NO_7_+H]^+^), MS^2^ fragmentation and chemical/biosynthetic logic, **23** is likely to be a 4-*O*-glucoside of menisdaurigenin, an isomer of menisdaurin (**11**) (**Fig. S10**).

*In vitro* enzymatic microsome assays were performed to attempt to identify the product of *Fs*CYP92B120 catalysed conversion of *p*-hydroxyphenylacetonitrile (**15**) (**Fig. S11**). However, no expected products could be identified, including oxo-menisdaurilide (**10**), the previously proposed substrate of *Fs*MS (**Fig. S11A**)^31^. Addition of purified *Fs*MS to *Fs*CYP92B120 microsomes with 4-hydroxyphenylacetonitrile (**15**) led to formation of menisdaurilide (**6**), which showed maximum formation over the 16 h time course, and menisdaurigenin (**12**), which accumulated in the first 3 h but disappeared across 16 h (**Fig. S11B**). This shows *Fs*CYP92B120 and *Fs*MS are responsible for menisdaurilide (**6**) formation, but reaction intermediates were unstable in our *in vitro* reaction or analytical conditions^33^.

Taken together, we propose that *Fs*CYP92B120 functions as a 4-hydroxyphenylacetonitrile oxidase, catalysing the oxidation of 4-hydroxyphenylacetonitrile (**15**) to produce the reactive intermediate oxo-menisdaurigenin (**17**). In *N. benthamiana*, in the absence of *Fs*MS, endogenous reductases act sequentially on **17** to first reduce the enone alkene moiety (i.e. Michael reduction) and then reduce the carbonyl to yield menisdaurin E isomers (**22**), which can non-enzymatically hydrolyse and cyclise into dihydromenisdaurilide (**21**). When *Fs*MS is present, it competes with the endogenous reductases for **17**, reducing it into menisdaurigenin (**12**), which then yields menisdaurilide (**6**) through hydrolysis and cyclisation.

### Menisdaurigenin is an intermediate of menisdaurilide biosynthesis

To further resolve the final steps of menisdaurilide biosynthesis *in planta*, including the identity of any short-lived intermediates, we conducted time-course experiments of *N. benthamiana* infiltrations, feeding 4-hydroxyphenylacetonitrile (**15**) to infiltrated plants (**Fig. S12** and **S13**). We found that with *Fs*CYP92B120 and *Fs*MS present, menisdaurilide (**6**) showed accumulation at 24 hours. Similarly, in the *Fs*CYP92B120 only sample we saw accumulation of oxo-menisdaurilide (**10**) over the same period. This contrasts with menisdaurigenin (**12**) which accumulates quickly in the *Fs*CYP92B120 and *Fs*MS sample, reaching a maximum at 3 hours, and then decreasing. Remarkably, we were able to identify a peak corresponding to the mass of the proposed *Fs*CYP92B120 product oxo-menisdaurigenin (**17**) (*m/z* = 150.0548, [C_8_H_7_NO_2_+H]^+^) which, in the *Fs*CYP92B120 only sample, accumulated rapidly, reaching a maximum at 1 h before decreasing by over the next 2 h. The peak is also present, though lower, in samples with both *Fs*CYP92B120 and *Fs*MS, also showing a maximum at 1 h.

Based on this time-course data (**Fig. S12** and **S13**) it appears likely that *Fs*CYP92B120 catalyses rapid formation of **17**, which is a substrate of *Fs*MS, forming menisdaurigenin (**12**) within 3 hours. The formation of menisdaurilide (**6**) from menisdaurigenin (**12**) is much slower, occurring over the course of 24 h, which matches the slow decline of menisdaurigenin (**12**) over this period. A similar slow accumulation of oxo-menisdaurilide (**10**) over 24 h in the absence of *Fs*MS is also observed.

Taken together, we have been able to elucidate the biosynthetic pathway from tyrosine (**5**) to menisdaurilide (**6**), which proceeds unexpectedly via nitrile intermediates (**Fig. S14**). We have also been able to identify multiple shunt products produced via endogenous enzymes in our expression systems, alongside the identification of three new cytochrome P450s and the activity revision of the previously described menisdaurilide synthase (*Fs*MS) into menisdaurigenin synthase. As a validation of this discovery, we reconstructed the pathway in *Saccharomyces cerevisiae*, using galactose inducible episomal plasmids. We transformed yeast with four pathway genes (*Fs*CYP79DC12, *Fs*CYP81BD45, *Fs*CYP92B120 and *Fs*MS) alongside a *F. suffruticosa* cytochrome P450 reductase (*Fs*CPR). When all pathway genes were present, aquilegiolide (**6b**) was produced (**Fig. S15**).

### *Fs*CYP92B120 and *Fs*MS are together responsible for the divergence of cyanogenic glucoside and menisdaurilide biosynthesis

The menisdaurilide (**6**) biosynthesis pathway shares steps with the dhurrin (**13**) cyanogenic glucoside pathway (**Fig. 1B** and **Fig. S14**). To further investigate the biochemical connection between these pathways, we tested whether enzymes from each pathway were functionally interchangeable (**Fig. 4G, Fig. S16**). The dhurrin pathway in *Sorghum bicolor* is catalysed by two cytochrome P450s: *Sb*CYP71E1, which converts tyrosine (**5**) into aldoximes (**14**), and the bifunctional *Sb*CYP79A1, which catalyses sequential dehydration and hydroxylation of the aldoxime to form *p*-hydroxymandelonitrile (**16**). Glucosylation of *p*-hydroxymandelonitrile (**16**), catalysed by *Sb*UGT85B1, converts it to dhurrin (**13**); this activity also appears to be endogenous to *N. benthamiana*. The glucosyl moiety in dhurrin protects it from degrading to *p-*hydroxybenzaldehyde and cyanide. We found that *Fs*CYP79DC12 and *Fs*CYP81BD45 could substitute *Sb*CYP79A1 and *Sb*CYP71E1 in the formation of dhurrin (**Fig. 4G**), indicating that *Fs*CYP81BD45 has some bifunctional activity like *Sb*CYP79A1. Addition of *Fs*CYP92B120 boosted dhurrin (**13**) formation further, indicating this enzyme can catalyse hydroxylation of 4-hydroxyphenylacetonitrile (**15**) to yield *p*-hydroxymandelonitrile (**16**). Substitution of *Fs*CYP81BD45 and *Fs*CYP92B120 with bifunctional *Sb*CYP71E1 in the presence of *Fs*CYP79DC12 and *Fs*MS did not lead to menisdaurilide (**6**) formation, indicating that *Fs*CYP92B120 activity is crucial for menisdaurilide formation. Furthermore, when *Fs*CYP79DC12 and *Fs*CYP81BD45 are co-infiltrated in *N. benthamiana*, dhurrin (**13**) accumulates, though addition of *Fs*CYP92B120 boosts production further (**Fig. S16**). These results demonstrate that the combined activities of *Fs*CYP92B120 and *Fs*MS lead to the bifurcation of the menisdaurilide (**6**) and dhurrin (**13**) pathways (**Fig. S14**).

### *De novo* biosynthesis of *Securinega* alkaloids

The entry point to the *Securinega* alkaloids is a Mannich reaction between the iminium electrophile Δ¹-piperideine (**4**) and the butenolide nucleophile menisdaurilide (**6**). This reaction can occur non-enzymatically under aqueous conditions, to yield the neosecurinanes^30,31^. The nonsymmetric formation of Δ¹-piperideine (**4**) from lysine (**3**) in *F. suffruticosa* is mediated by *Fs*OLADO^18^. Therefore, we expected that the menisdaurilide biosynthetic pathway was the final piece required for *de novo* biosynthesis of *Securinega* alkaloids. Indeed, when the four-gene menisdaurilide pathway was co-infiltrated with *Fs*OLADO in *N. benthamiana* we observed *de novo* formation of the fused-ring neosecurinane alkaloids virosine A (**8**) and securinol A (**9**) (**Fig. 5A** and **Fig. S17**). We also observed the formation of a direct non-fused-ring Mannich product that we characterised and named flueggeanine (**24**) (**Fig. 5A** and **Fig. S17**); this was confirmed to be a *F. suffruticosa* natural product (**Fig. S18**). We also demonstrated *in vitro* formation of *Securinega* alkaloids using purified enzymes (*Fs*MS and *Fs*OLADO) and yeast derived microsomes (3x cytochrome P450s) with lysine (**3**) and tyrosine (**5**) as substrates, also observing formation of securinol A (**9**), virosine A (**8**) and flueggeanine (**24**) (**Fig. 5A**).

**Figure 5.**
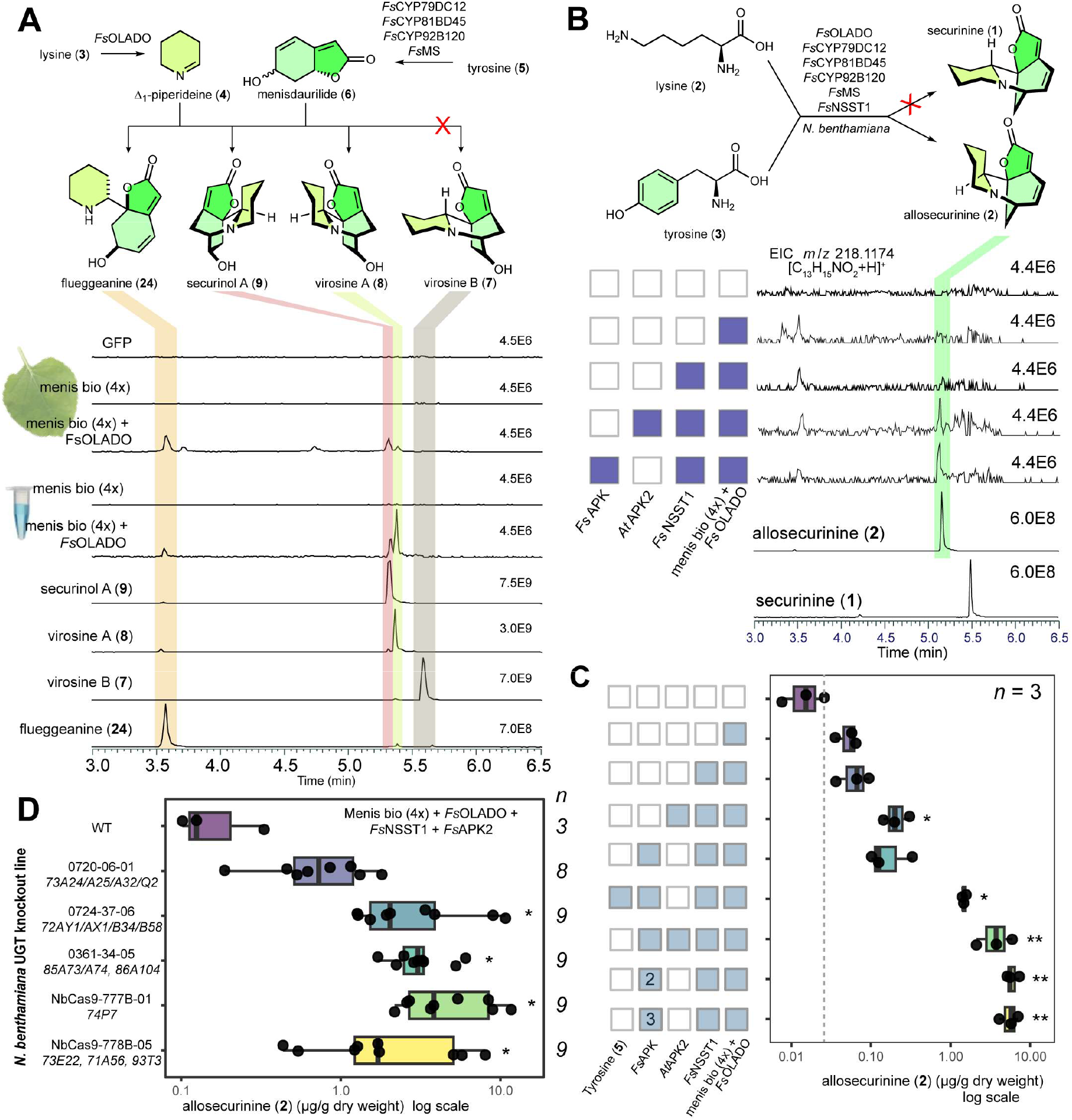
*De novo* biosynthesis of *Securinega* alkaloids. (**A**) Formation of neosecurinanes (**7**, **8** and **9**) and flueggeanine (**24**) in *N. benthamiana* (leaf) and *in vitro* (tube) from lysine (**3**) and tyrosine (**5**) with menisdaurilde biosynthesis genes (menis bio (4x)) and *Fs*OLADO (EICs *m*/*z* 236.1280). (**B**) Formation of allosecurinine (**2**) in *N. benthamiana* leaves by adding *Fs*NSST1 and APKs to neosecurinane genes (panel **A**) (EICs *m*/*z* 218.1174). (**C**) Allosecurinine (**2**) accumulation with pathway boosting strategies. *Fs*APK numbers refer to higher infiltration dosage. Dotted line shows average background peak area; stars show significant differences to control (GFP, row 1). (**D**) Allosecurinine (**2**) accumulation with *N. benthamiana* UGT mutant lines. UGTs knocked out are in italics, stars show significant differences to WT. See **Fig. S17** for MS^2^ validation of peaks; statistical tests are Welch ANOVA + Games-Howell.

Next, we sought to gain access to the securinane alkaloids, such as securinine (**1**) and allosecurinine (**2**). To do this we used a sulfotransferase *Fs*NSST1 recently reported to catalyse the conversion of neosecurinanes to securinanes^31^. Addition of just *Fs*NSST1 to the five-enzyme neosecurinane pathway in *N. benthamiana* did not result in significant formation of securinane alkaloids (**2**) (**Fig. 5B, 5C**). To enhance *Fs*NSST1 activity, we introduced adenosine-5′-phosphosulfate kinases (APKs) from *Arabidopsis thaliana* (*At*APK2) or *F. suffruticosa* (*Fs*APK2), which boost production of the essential sulfotransferase cofactor 3′-phosphoadenosine-5′-phosphosulfate^40^. Addition of *At*APK2 to the six-enzyme pathway resulted in *de novo* production of allosecurinine (**2**) significantly (14-fold) above background (**Fig. 5C** and **Fig. S17C**). Supplementation of tyrosine (**5**) or higher doses of APK2s led to further 100–400-fold increases of allosecurinine (**2**) accumulation (**Fig. 5C**). No formation of securinine (**1**) was observed (**Fig. 5A**).

Based on the accumulation of glucosylated side-products in our *N. benthamiana* reconstruction caused by endogenous glucosyltransferases, we anticipated that plants with specific UGTs knocked out may accumulate fewer glucoside shunt products and therefore enable greater flux to the alkaloids^41^. We examined *Securinega* alkaloid production in five different *N. benthamiana* UGT-knock out lines: four lines showed significant increases of allosecurinine (**2**) formation compared to wild-type, with the best performing mutant showing an average 30-fold increase in allosecurinine (**2**) production compared to the wild-type (**Fig. 5D**).

## Discussion

Complex alkaloids are typically formed through the Mannich condensation of an amino acid derived imine electrophile and a nucleophile^1^. In the biosynthesis of *Securinega* alkaloids such as securinine (**1**) and allosecurinine (**2**), the imine electrophile is Δ¹-piperideine^18^ which is also the precursor to diverse alkaloid families including lupin and *Lycopodium* alkaloids. The unique feature of *Securinega* alkaloid origin is the nucleophile, which was determined, through biomimetic syntheses^27^ and natural product isolation^25,26^, to be the butenolide menisdaurilide (**6**). Whilst labelling experiments showed menisdaurilide (**6**) was expected to derive from tyrosine (**5**), its specific biosynthetic origin was unknown, preventing biocatalytic entry to the *Securinega* alkaloid pathways.

We have determined that menisdaurilide (**6**) is derived from tyrosine (**5**) via a cyanogenic glucoside-like pathway involving aldoxime and nitrile intermediates that lose their nitrogen-containing functional groups at the final step. Previous evidence supported two key aspects of the pathway: the non-enzymatic formation of menisdaurilide (**6**) from a nitrile precursor (menisdaurin (**11**))^33^ and the origin of nitrile glucosides like **11** from cyanogenic glucoside-like pathways^36^. Here we connected these observations to *F. suffruticosa* alkaloid biosynthesis, enabling pathway elucidation from tyrosine (**5**) to menisdaurilide (**6**).

The first steps of the menisdaurilide pathway, formation of aldoxime and then nitrile from an amino acid, are common across multiple plant pathways, including camalexin^42^, auxins^43^ and cyanogenic glucosides. We show that a similar pathway contributes to the formation of nitriles, butenolides and, most remarkably, complex alkaloids. This connection extends beyond shared chemical intermediates. Aldoxime formation in *F. suffruticosa* is catalysed by a CYP79 family enzyme, typical in angiosperm cyanogenic glucoside formation^44^. Moreover, genes from cyanogenic glucoside biosynthesis are often found in biosynthetic gene clusters^45^: in the *F. suffruticosa* genome the menisdaurilide genes do not form a complete biosynthetic gene cluster but those responsible for the first two pathway steps are adjacent (**Fig. S2**).

We demonstrated a biochemical connection between dhurrin (**13**) and menisdaurilide biosynthesis. *F. suffruticosa* cytochrome P450s (together with an endogenous *N. benthamiana* UGT) produce dhurrin (**Fig. S16**). Due to the enzyme bifunctionality, dhurrin can be formed by *Fs*CYP79DC12 with either *Fs*CYP81BD45 or *Fs*CYP92B120. Crucially, *Fs*CYP92B120 oxidises *p*-hydroxyphenylacetonitrile (**15**) into either *p*-hydroxymendelonitrile (**16**), the dhurrin precursor, or oxo-menisdaurigenin (**17**), the menisdaurilide precursor, driving the biochemical divergence of these pathways. In nature, dhurrin (**13**) and menisdaurin (**11**) co-occur in *Tiquilia* species^36^, whilst tyrosine derived cyanogenic glucosides and menisdaurilide are both found in Phyllanthaceae^46–48^. Cyanogenic glucosides have evolved independently multiple times^44^ and Phyllanthaceae represents another example of this, but with the core pathway diversifying towards nitrile glucosides, butenolides and alkaloids.

*Nicotiana benthamiana* is an invaluable tool for reconstructing plant natural product biosynthesis^49^ but its endogenous UGTs frequently glucosylate xenochemicals, probably targeting them for transport into the vacuole^50^. We observed at least four *N. benthamiana* UGTs catalysed conversions including dhurrin (**13**) formation from *p*-hydroxymandelonitrile (**16**), possibly through the action of *Nb*UGT85A73^51^. Infiltrating *N. benthamiana* UGT knock-out lines^41^ significantly increased allosecurinine formation compared to wild-type in four out of five lines, suggesting multiple UGTs were responsible for restricting flux through the pathway.

Additionally, multiple shunt products arose from the sequential reductions of oxo-menisdaurigenin (**17**): a 1,4-Michael reduction followed by a 1,2-carbonyl reduction. This forms menisdaurin E (**22**) and the hydrolysis product dihydromenisdaurilide (**21**). Similar two-stage enone reductions catalysed by endogenous *N. benthamiana* enzymes have been observed in artemisinin^52^ and digitoxin biosynthesis^53^. Knocking-out these promiscuous reductases could further boost pathway flux^54^.

The reductase *Fs*MS was previously reported as menisdaurilide synthase, proposed to reduce oxo-menisdaurilide (**10**) to menisdaurilide (**6**)^31^. However, when co-expressed with *Fs*CYP92B120, *Fs*MS competes with the endogenous reductases for oxo-menisdaurigenin (**17**), decreasing shunt products (**21** and **22**) whilst increasing menisdaurigenin (**12**) and its glucosylated shunt **23**. Furthermore, the butenolides oxo-menisdaurilide (**10**) and menisdaurilide (**6**) only accumulate slowly (**Fig. S13**). Thus, *Fs*MS preferentially reduces oxo-menisdaurigenin (**17**) to menisdaurigenin (**12**) in our system. The observation of menisdaurin (**11**) in *Flueggea* species further indicates menisdaurigenin (**12**) is an *in planta* intermediate, leading us to reconsider *Fs*MS as a menisdaurigenin synthase. In our pathway reconstructions *in planta*, *in vitro* and in yeast the conversion of menisdaurigenin (**12**) to menisdaurilide or aquilegiolide (**6**) occurs without the addition of a specific enzyme to control this step. However, time course experiments in *N. benthamiana* suggest that this conversion is relatively slow, creating a bottleneck that an enzyme might accelerate in nature.

We achieved *de novo* formation of *Securinega* alkaloids, forming the neosecurinanes virosine A (**8**) and securinol A (**9**), and, with *Fs*NSST1 and helper APK2s, allosecurinine (**2**). We did not observe securinine (**1**), likely due to the absence of its neosecurinane precursor, virosine B (**7**), a minor product of the non-enzymatic Mannich reaction^30^. This contrasts with the abundance of securinine (**1**) in *F. suffruticosa* (**Fig. S18**). This discrepancy might be resolved by discovery of a coupling enzyme that controls the selectivity of the Mannich reaction, similar to the carbonic anhydrase-like enzymes that control the stereoselective Mannich reaction in *Lycopodium* alkaloid biosythesis^20^. Alternatively, an oxidation-reduction epimerisation reaction may convert allosecurinine (**2**) into securinine (**1**), as seen in benzylisoquinoline^55,56^ and monoterpene indole alkaloid biosynthesis^57,58^.

Elucidating plant biosynthesis pathways enables heterologous production of valuable bioactive compounds. It also reveals how metabolic diversity arises in plants, metabolically, enzymatically, and through evolution. The discovery of menisdaurilide biosynthesis unlocks *Securinega* alkaloids for engineering, highlights the metabolic and enzymatic interplay between different natural product classes and demonstrates how pathways may be repurposed to generate chemical diversity.

## Methods

### Candidate gene screening, cloning and expression vector construction

Gene co-expression analysis was performed by using the *Fs*OLADO gene as the bait gene and the published transcriptome data^18^. Transcripts per million (TPM) values for each gene were obtained, log transformed [log_2_(TPM+1)] and a Pearson’s correlation coeffcient (PCC) with *Fs*OLADO (gene1864) was calculated. Genes which had PCC(*Fs*OLADO) above 0.75 were investigated, and those annotated as CYP450s, glucosyltransferases, reductases, *β*-glucosidases were selected as candidates.

To obtain cDNA for cloning candidates, *Flueggea suffruticosa* young leaves were taken from branches with active growth and immediately flash frozen in liquid nitrogen. Material was ground into a fine powder using a mortar and pestle under liquid nitrogen. Ground tissue (∼50 mg) was used for total RNA extraction using a RNeasy Plant Mini Kit (QIAGEN), following kit instructions for the RLT buffer homogenisation method. Extracted RNA was then treated using a RNA Clean & Concentrator-5 kit (ZYMO RESEARCH) following kit instructions. cDNA was synthesised using extracted and cleaned RNA (5 µL) with Invitrogen™ SuperScript™ IV First-Strand Synthesis System, following kit instructions.

Primers were designed in Geneious Prime and synthesized by Merck. For the Goldenbraid method to clone genes of interest into the desired vector, primers containing corresponding overhangs were also synthesized (**Table S2**). PCR reactions were performed by using Platinum SuperFi^TM^ polymerase enzyme (Thermo Fisher Scientific), and amplified genes were identified on an agarose gel and purified using the Monarch^®^ PCR & DNA Cleanup Kit (New England Biolabs). The blunt end fragments were inserted into the pJET clone vector using the CloneJET PCR cloning Kit (Thermo Fisher Scientific). Gene domestication and expression vector construction were conducted by following the Goldenbraid cloning protocol^59^. The constructs were transformed into Stellar competent cells (Takara Bio) and confirmed by whole plasmid sequencing (Plasmidsaurus). For *E. coli* expression, a pPGN-K expression vector was used^60^; for *N. benthamiana* expression, the vector pDGB3-α1 was used, and for yeast expression, domesticated pAV series vectors (pRS423 backbone) were used.

### Transient expression in *Nicotiana benthamiana*

Expression constructs (pDGB3-α1-GOI) with confirmed sequence content were transformed into *Agrobacterium* GV3101 chemically competent cells, which then were spread on LB agar plates (50 μg/mL kanamycin, 25 μg/mL gentamicin, 10 μg/mL rifampicin) and incubated (28 °C, 2-3 d). Colonies appearing on the LB plates were verified by colony PCR (GoTaq green master mix, Promega) and, if positive, inoculated into LB liquid media with the corresponding antibiotics. Cultures were incubated (28 °C, 200 rpm, 24 h), pelleted by centrifugation (20 min, 3500 rpm) and resuspended in infiltration buffer (10 mM MES pH 5.5, 10 mM MgCl_2_, 100 μM acetosyringone) to reach OD 600 = 0.5. *N. benthamiana* plants were grown in F2+S soil in a growth room before and after infiltration (25°C, 16 : 8 h, light : dark). Cultures were infiltrated into the abaxial side of 5 weeks old plant leaves using a 1 mL syringe. Two leaves per plant and three plant replicates per group were used. A pHREAC-GFP containing culture was used as the negative control. Leaf samples were harvested after 5 d of infiltration. If necessary, substrates were supplemented after 4 d of infiltration and samples were harvested after two more days. For time-course experiments the sampling time is recorded in the associated figures. For leaf disc incubation experiments, leaf discs (8 mm diameter) were collected after 4 d of infiltration and positioned epidermis facing up into 24-well plates with an incubation buffer (500 μL, 50 mM HEPES, pH 7.5). Virosine B (**7**) was then added into the buffer at a final concentration of 50 μM. Leaf discs were incubated at the growth room for 24 h and harvested for further analysis.

### *In vitro* enzymatic assays

To characterize the function of *Fs*UGT85A242, *in vitro* assays were performed in a reaction mixture containing HEPES buffer (50 mM, pH 7.5), aldoximes (**14**) (2 mM), purified *Fs*UGT85A242 protein (10 ng/μL), dithiothreitol (DTT) (2 mM), NaCl (50 mM), and UDP-glucose (UDPG) (5 mM), with the final volume adjusted to 100 μL with water. For cytochrome P450 assays, reactions were carried out in 20 mM sodium phosphate buffer (20 mM, pH 7.0 containing microsomal fraction (10 μL), purified *Fs*MS protein (10 ng/μL, when required), purified *Fs*OLADO (10 ng/μL, when required), NADPH (reduced nicotinamide adenine dinucleotide phosphate, 5 mM), and L-tyrosine (**5**) (1 mM), *L*-lysine (**3**) (1 mM) (when required), with the final volume adjusted to 100 μL. All reaction mixtures were either incubated at 30 °C overnight or harvested at different time points, as noted in the figures. Heat-denatured enzymes (95 °C for 10 min) were used as negative controls.

### LC-MS sample preparation and detection

For *N. benthamiana* samples, three leaf discs were combined into one tube and flash frozen in liquid nitrogen. Samples were homogenized to fine powder using the Tissuelyser (30 s, 30 Hz) whilst frozen and then resuspended in 80% v/v methanol (500 μL). Tubes were vortexed at room temperature (1500 rpm for 1 h), then centrifuged (15,000 rpm, 25 min) to obtain the supernatant. The *in vitro* enzyme assays were quenched with methanol (100 μL). The tube was vortexed vigorously (10 s) and then centrifuged (15,000 rpm, 25 min) to obtain the supernatant. For engineered yeast metabolic analysis, induced yeast cultures (100 μL) were sampled and yeast cells were harvested by centrifugation (10,000 rpm, 5 min). Methanol (100 μL) was added to the growth media and centrifuged to obtain the supernatant (15,000 rpm, 25 min). The supernatant was transferred to LC-MS glass vials for further analysis.

Liquid chromatography was performed using a Waters Acquity I-Class UPLC system and the column was a Waters UPLC BEN C18 column (130 Å, 1.7 µm, 2.1 mm X 50 mm) which was compatible for analyses conducted in high pH conditions. The column temperature was maintained at 60 °C. Sample injection volume was 2 μL. The mobile phase consisted of (A) 10 mM ammonium bicarbonate at pH 10.2, adjusted with ammonium hydroxide, and (B) methanol. The flow rate was set at 0.5 mL/min. Gradient %B: 0 min 2%; 4 min 40%; 4.5 min 100%; 5.5 min, 100%; 6.1 min 2%; 8 min 2%. For time course experiments the mobile phases consisted of (A) 0.1% formic acid in water and (B) 0.1 formic acid in methanol. The flow rate was 0.45 mL/min. Gradient %B: 0 min 2%; 6.5 min 100%; 8 min 100%; 8.5 min 2%; 10 min 2%. Mass spectrometric (MS) was performed on a Thermo Fusion Orbitrap mass spectrometer with an Atmospheric Pressure Chemical Ionization (APCI) source, operating in positive ion mode. Mass spectral data were acquired over the *m/z* range of 50 to 500. Data was analysed on Freestyle software.

### Assessment of allosecurinine formation

After infiltration, transformed *N. benthamiana* leaf samples were harvested after 5 d of infiltration and freeze-fried. Samples (10.8 mg) were extracted into 80% v/v methanol (500 μL) and analysed by LC-MS as described above. Allosecurinine (**2**) peak areas were converted to concentrations using an external standard curve and expressed as µg/g dry weight. Comparisons were analyzed on log_10_-transformed values by Welch’s ANOVA, followed by Games-Howell post hoc pairwise comparisons, appropriate for unequal variances and sample sizes. For comparison, an average background peak area was determined using the average peak areas in the no enzyme controls across a retention time range of equal width to allosecurinine (**2**) peak but adjacent and non overlapping to the retention time of the standard. Statistical significance was set at *p* < 0.05 (* = *p* < 0.05, ** = *p* < 0.01). All analyses were performed in *R* (v4.x) using the rstatix and car packages.

### GC-MS sample preparation and test

*In vitro* enzymatic assays were quenched with ethyl acetate (100 μL). The tube was vortexed (RT, 10 min) and left to settle (10 min). The top ethyl acetate layer was transferred into the GC-MS glass vial for further analysis. Agilent 7890A gas chromatograph (Agilent Technologies) interfaced to an Agilent 5975C MSD (Agilent Technologies) was used for the GC-MS analyses of all the samples via direct injection. The separation was performed in splitless mode on a Zebron ZB5-HT-INFERNO column (length: 30 m; diameter: 250 μm; Phenomenex) interfaced with auxiliary transfer line (1.5 m, 150 μm). Helium carrier gas was used at 1.2 mL/min constant flow and the transfer line temperature was set to 280 °C. The oven temperature was set at 80 °C for 5 min and then increased to 110 °C at a rate of 2.5 °C/min followed by increase to 280 °C at a rate of 10 °C/min. 5 µL of each sample were injected. Ethyl acetate was used to wash the needle between each injection. The mass scan range was 40–500 *m/z* on normal scanning mode.

### Chemicals used in the study

Menisdaurin (**11**) was purchased from MedChemExpress. L-Tyrosine (**5**), 4-hydroxyphenylacetonitrile (**15**), and L-lysine (**3**) were purchased from Merck (Sigma-Aldrich). All the other chemicals were synthesized in our lab and the methods and NMR data were summarized in **Supplementary information.**

### Recombinant protein expression and purification

Sequence verified plasmids (pPGN-K-*Fs*MS, pPGN-K-*Fs*UGT85A242) were used to transform SoluBL21 *E. coli* expression competent cells by heat-shock method. A single colony was selected from the overnight plates and inoculated into individual Erlenmeyer flasks (100 mL volume) containing LB broth (20 mL with 50 μg/mL kanamycin). The flasks were incubated overnight (37 °C, 220 rpm). Overnight culture (10 mL) was added to each flask (2.5 L volume) containing 2x YT (Yeast Extract Tryptone) media (1 L with 50 μg/mL kanamycin). The culture was incubated (37 °C, 220 RPM) until 0.5≥OD600. The cultures were cooled to 18 °C and then IPTG (Isopropyl β-D-1-thiogalactopyranoside, 1 mM) was added into each culture to induce recombinant protein production. Flasks were then incubated overnight (16–20 h, 18 °C, 200 rpm). Cultures were harvested using the through centrifugation (Sorvall LYNX 4000 centrifuge (Thermo Fisher Scientific), 4 °C, 5000 xg, 30 min). The supernatant was discarded once clear, and pellets were resuspended in Lysis Buffer [50 mM Tris-HCL (pH 8.0), 50 mM glycine, 5% v/v glycerol, 0.5 M NaCl, 20 mM imidazole, 1 x EDTA-free protease inhibitor tablet, and 0.2 mg/mL lysozyme], and incubated (30 min, 4 °C, with gentle rotation)). For efficient lysis, each *E. coli* suspension was loaded onto a pre-cooled Cell/French Press at 25 Kpsi. The suspensions were then centrifuged (4 °C, 20 min, 35000 xg) to remove cell debris. For benchtop purification of recombinant proteins, Ni-NTA resin (500 μL) was washed in water and then binding buffer [50 mM Tris-HCl (pH 8.0), 50 mM glycine, 5% v/v glycerol, 0.5 M NaCl, 20 mM imidazole] then added to clarified lysates. The mixture was incubated (4 °C, 1 h, with gentle rotation) to allow the His-tagged protein to bind the Ni-NTA. Samples were pelleted by centrifugation (1000 xg, 1 min) and the supernatant was gently poured away. The resin was washed in the Wash buffer [50 mM Tris-HCl (pH 8.0), 50 mM glycine, 5% v/v glycerol, 0.5 M NaCl, 40 mM imidazole] three times. Elution buffer [300 μL of 50 mM Tris-HCl (pH 8.0), 50 mM glycine, 5% v/v glycerol, 0.5 M NaCl, 500 mM imidazole] was used to elute the binding proteins off the resin. Buffer exchange was performed using a Vivaspin column to wash away the high imidazole and the protein was resuspended in storage buffer [50 mM HEPES (pH 7.5), 150 mM NaCl and 10% glycerol], and flash-frozen in liquid nitrogen.

### Yeast strain construction and microsome preparation

To express targeted genes in yeast, *S. cerevisiae* CEN.PK2-1C (MATα ura3-52; trp1-289; leu2-3,112; his3Δ 1; MAL2-8c; SUC2, purchased from EUROSCARF, accession no-30000B) was used in this study. To prepare yeast competent cells, a single colony of the CEN.PK strain was inoculated into YPD (Yeast Extract Peptone Dextrose) growth liquid media (5 mL) and incubated (30 °C, 200 rpm, overnight). We measured the OD600 of the overnight culture and then diluted it YPD liquid media (∼50 mL) to reach a final OD600 of 0.2. The culture was incubated (30 °C, 200 rpm) until OD600 reached 0.6 and then centrifuged (3500 rpm, 5 min) to harvest the pellet. The cell pellet was washed with water twice and then resuspended in sterile lithium acetate (LiAc, 100 mM). The competent cells were immediately used for transformation. The transformation mixture included: pAV plasmid (1 μg) containing single CYP450, pSEC-AaCPR (*Artemisia annua* cytochrome P450 reductase. 1 μg), ssDNA (single stranded DNA from salmon testes, 50 μL), yeast competent cells (90 μL) and PLATE buffer (900 μL, [50% polyethylene glycol (PEG) 2000, 0.1 M lithium acetate, 10 mM Tris-Cl, 1 mM EDTA, pH 8.0]). The mix was incubated (30 °C, 30 min) and heat shocked (42 °C, 45 min). The reaction was centrifuged (3500 rpm, 2 min) to collect cells and then cells were resuspended in YPD liquid media (500 μL) and incubated (30 °C, 200 rpm) for recovery. The recovered cells were plated onto their corresponding selective SD plates. For *Fs*CYP79DC12/ *Fs*CYP79DC13/ *Fs*CYP81BD45, SD-His+Leu plates were used; for *Fs*CYP92B120, SD-Ura+Leu plates were used. The plates were incubated (30 °C, 2–3 d) until colonies began to grow. Positive colonies were confirmed by colony PCR and used for subsequent analysis.

Confirmed colonies were inoculated into corresponding SD-drop-out liquid growth media containing 2% (w/v) glucose and incubated (30 °C, 220 rpm). The overnight culture was diluted to an OD600 of 0.05 in SD-drop-out liquid medium supplemented with 2% glucose and incubated (16 h). The yeast pellets were harvested and resuspended in SD-drop-out liquid medium supplemented with 2% galactose and then cultured (24 h, 30 °C, 220 rpm). The induced cells were harvested and resuspended and washed with TEK buffer (0.1 M KCl in TE buffer, 50 mM Tris, pH 7.4, 1 mM EDTA). After the wash, the supernatant was discarded and cells were resuspended in TES-B buffer (0.6 M sorbitol in TE buffer) and lysed by using the French pressure cell press. The lysate was then centrifuged (10,000 rpm, 30 min, 4 °C) and the supernatant was transferred to ultra-centrifuge tubes and centrifuged (35,000 rpm, 1 h, 4 °C). The microsome pellets were suspended in TEG buffer (20 % glycerol in TE buffer) and used for *in vitro* enzymatic assays.

### Yeast episomal transformation

InFusion cloning (Takara Bio) was performed to create the *Fs*CYP81BD45/*Fs*CYP92B120 dual gene cassette as follows. The *Fs*CYP92B120 gene in pAV126 the vector was linearised with the restriction enzyme XhoI via restriction digest (37 °C, 4 h). The fragment of *Fs*CYP81BD45 with restriction enzyme overhangs was amplified from its pAV123 vector with PCR (Platinum SuperFi^TM^, Thermo Fisher Scientific). PCR product cleanup was completed with the Monarch^®^ PCR & DNA Cleanup Kit (New England Biolabs). A 2:1 molar ratio of fragment to linearised vector was used in an InFusion reaction (10 µL, 50 °C, 40 min). A portion (5 µL) of this reaction was added to *E. coli* Stellar^TM^ cells (Takara Bio) (50 µL), followed by heat shock using a thermocycler (4 °C 25 min, 42 °C 45 s, 4 °C hold). The heat-shocked product was added to SOC media (500 µL), which was incubated (1 h, 37 °C, 200 rpm), and then plated on LB media (100 µg/ml ampicillin) and incubated overnight (37 °C). After this, colony PCR (GoTaq Green Master Mix, Promega) was used to genotype transformants. Transformants with the correct product size, as determined with agarose gel electrophoresis, were inoculated into LB medium (5 mL, with 100 µg/ml ampicillin) and grown overnight (37 °C, 200 rpm), followed by plasmid extraction with the ZymoPURE^TM^ Plasmid Miniprep Kit (Zymo Research). PCR product cleanup utilized the Monarch® PCR & DNA Cleanup Kit (New England Biolabs). For the InFusion reaction, a 2:1 molar ratio of fragment to linearised vector was employed (10 µL reaction volume, 50 °C, 40 min). Transformation was performed by adding a portion (5 µL) of the InFusion reaction to *E. coli* StellarTM cells (Takara Bio) (50 µL), followed by a heat shock protocol (4 °C for 25 min, 42 °C for 45 s, followed by a 4 °C hold). The resulting mixture was incubated in SOC media (500 µL, 1 h, 37 °C, 200 rpm). The cells were then plated on LB agar (with 100 µg ml^-1^ ampicillin) and incubated overnight (37 °C). Successful transformants were genotyped using colony PCR (GoTaq Green Master Mix, Promega). Transformants exhibiting the correct product size, confirmed via agarose gel electrophoresis, were inoculated into LB media (5 mL with 100 µg mL^-1^ ampicillin) and grown overnight (37 °C and 200 rpm). Finally, plasmid extraction was carried out using the ZymoPURETM Plasmid Miniprep Kit (Zymo Research). *Fs*MS was cloned into pBEVY-T vector^61^ through restriction digestion with EcoRI and KpnI, with further steps as described above.

Episomal expression was achieved through plasmid transformation with the lithium acetate method^62^. Plasmids used for transformation were pAV123-*Fs*CYP79DC12, pAV125-*Fs*CPR, pAV126-*Fs*CYP81BD45/*Fs*CYP92B120, pBEVY-T-*Fs*MS. Transformation culture (150 µL) was plated onto SC-glucose media with AAs (amino acids) for selection and incubated (72 h, 30 °C). After validation with colony PCR (GoTaq Green Master Mix, Promega), liquid cultures (1 mL) were grown (30 °C, 700 rpm) in SC-glucose media with AAs in the 24-well growth plate. To induce gene expression, yeast cultures were grown (24 h) and pelleted through centrifugation (3000 *x g* for 10 min) and resuspended in SC media (1 mL) with 2% wt/vol galactose, and appropriate AAs. These cultures were incubated for 24 h or 72 h at 700 rpm and 30 °C prior to metabolite analysis.

## Availability of data and materials

Sequences of coding sequences of genes will be available on Genbank at publication.

## Acknowledgements

We thank Dr Nicola Patron for providing the UGT-knockout *N. benthamiana* seeds and Dr Elizabeth Rylott and Dr Giuliano Pechar for providing *Sorghum bicolor* plant material and plasmids harbouring dhurrin biosynthetic pathway genes. We thank Prof David Nelson for providing the P450 names and Prof Michael Court and the UGT Nomenclature Committee Chair for the UGT name. We thank the Centre for Excellence in Mass Spectrometry for access to mass spectrometry equipment, and the Biosciences Technology Facility for technical support, especially Tony Larson and Martina Conte. We also gratefully acknowledge the University of York’s Chemistry Analytical Facility and Horticultural facility for their support & assistance in this work. We acknowledge valuable discussions with Prof Sang-Gyu Kim and his team including Taein Kim. The plasmid pHREAC was a gift from George Lomonossoff (Addgene plasmid #134908). BRL acknowledges support from UKRI (MR/S01862X/1) and BBSRC (BB/Y003586/1).

## Supplementary Data

Supplementary Information

Supplementary Figures (Figs. S1 to S18)

Supplementary Tables (Table S1 and S2)

Synthetic Chemistry

## Author contributions

ZJ carried out the gene cloning, enzyme purification and characterization *in vitro* and in *N. benthamiana* with the assistance of IA. LJNW and ORH performed chemical synthesis. CH and ZJ constructed engineered yeast stains. WPU assisted in the chemical synthesis and interpretation of NMR data. GG assisted in the protein expression and purification analysis. BRL initiated the project and assisted in the interpretation of data. ZJ and BRL wrote the manuscript with input from all authors.

